# An evolutionary shift to prioritizing mating over care is associated with consistently high androgen levels in male threespine stickleback

**DOI:** 10.1101/2025.04.11.648435

**Authors:** Meghan F. Maciejewski, Alison M. Bell

## Abstract

Steroid hormones play a role in regulating social behaviors in vertebrates, but how they mediate the evolution of these traits remains an open question. Here, we use liquid chromatography-mass spectrometry (LC-MS/MS) to quantify a panel of steroids in breeding males of two recently diverged populations of threespine stickleback. The common ecotype provides paternal care, while the white ecotype has evolutionarily lost paternal care. Hormone levels were quantified in both ecotypes at three reproductive stages: (1) after completing a nest, (2) soon after mating, when commons start to provide care and whites disperse the embryos, and (3) four days after mating, when commons are performing parental care and no longer courting females while whites are not providing care and are courting females. Androgens declined in the common ecotype as they began providing care but remained elevated in the white ecotype across stages, possibly to maintain the production of “spiggin,” the androgen-dependent glue males use to construct nests. Progestogen levels were low in whites and were lowest in commons after mating, suggesting an antagonistic relationship between progestogens and sexual behavior. Both ecotypes showed elevated glucocorticoids after mating, suggesting the stress axis may not have diverged between ecotypes. Altogether, these results provide evidence that the ecotypes regulate steroid levels differently to support the ways they balance mating and parental effort. Our data suggest a variety of mechanisms by which steroid signaling and regulation can change during the early stages of divergence between behaviorally distinct populations.

## 1. INTRODUCTION

Steroid hormones coordinate the expression of many traits and regulate behavior, physiology, and morphology in vertebrates. Steroids are particularly important for reproduction, as they coordinate the timing of gamete production, expression of secondary sexual characteristics, and behaviors like courtship and nesting with external cues such as photoperiod and the presence of potential mates (Adkins-Regan, 2005). The Challenge Hypothesis, first described in birds (Wingfield et al., 1990) and later extended to other vertebrate taxa (Hirschenhauser and Oliveira, 2006), proposes a central role for androgens in regulating male reproductive behavior and trade-offs between aggression and care. The role of androgens in coordinating suites of traits in male vertebrates is conserved (Hau 2007), but the components of androgen signaling, i.e., “endocrine building blocks,” can show substantial variation (Rosvall, 2022). As products of complicated endocrine cascades involving the brain, pituitary, and testes (hypothalamic-pituitary-gonadal (HPG) axis), androgen production and signaling can vary in a multitude of ways, including expression of steroidogenic enzymes, binding globulins, co-factors, receptor abundance, and receptor affinity (Rosvall, 2022).

Because steroids play a central role in regulating suites of traits like spermatogenesis, aggression, care, and secondary sexual characteristics, mechanisms regulating their production and signaling can likely facilitate or constrain the evolution of these traits (Hau, 2007; Ketterson et al., 2009), yet how hormones mediate the evolution of behavior and other phenotypes remains an open question (Cox et al., 2016). Determining if and how hormone levels vary among divergent groups is an important first step toward understanding how steroid-mediated traits evolve. Once steroid levels have been established, one can begin to narrow in on how these differences occur. Intraspecific comparisons of hormone levels in closely related populations (e.g., Bergeon Burns et al., 2013; Enbody et al., 2018; Graham et al., 2018; Kitano et al., 2010, 2011; Kitano and Lema, 2013; Miles et al., 2018) can provide particularly powerful insight into the hormonal mechanisms underlying behavioral evolution, as there has been less time for differences confounded with the behavior of interest to emerge (Jourjine and Hoekstra, 2021).

Here, we compare whole-body levels of steroids between two closely related ecotypes of threespine stickleback (*Gasterosteus aculeatus*) with dramatic differences in their reproductive behavior. Typically, male stickleback are solely responsible for providing paternal care that is necessary for offspring survival (van Iersel, 1953). A few days after fertilization, males transition from the “sexual phase,” in which they build nests and court females, into the “parental phase,” during which they exclusively care for their offspring (van Iersel, 1953; Jamieson et al., 1992). However, an atypical ecotype, known as the “white” ecotype due to its nuptial coloration, has recently diverged from the typical “common” caregiving ecotype (Samuk, 2016) and evolutionarily lost paternal care (Blouw and Hagen, 1990; Blouw, 1996). Whites spit their offspring out of the nest within minutes of fertilization (Jamieson et al., 1992; Blouw, 1996). Unburdened by the constraints of parental care, male whites then resume courting females, allowing them to invest nearly twice as much time in seeking new mates than commons (Jamieson et al., 1992; Blouw, 1996). These two ecotypes differ not only in parental behavior and coloration (Blouw and Hagen, 1990; Blouw, 1996) but also in a suite of other traits, including nest architecture (Behrens et al., 2024), courtship behavior (Behrens et al., 2024), and juvenile social behavior (Neumann and Bell, 2023). Given the important role androgens play in coordinating male reproductive physiology and behavior in vertebrates, including stickleback (Mayer et al., 2004), we hypothesized that the two ecotypes exhibit different androgen profiles resulting from differences in how they regulate androgen levels throughout the breeding season.

Previous work has demonstrated that androgen levels, particularly 11-ketotestosterone (11-KT), the major androgen in stickleback (Mayer et al., 2004), peak when males are attracting mates and drop when males are providing care (Páll et al., 2002a). Therefore, we predicted that whites, which do not provide paternal care, do not experience this drop in androgens and instead maintain high levels of androgens throughout the breeding season. We predicted that commons experience a drop in 11-KT when they begin to parent, consistent with previous work (Páll et al., 2002a). Although the drop in 11-KT has been shown to coincide with a rise in parental fanning behavior and a decline in courtship (Páll et al., 2002a), the administration of androgens does not affect the performance of these behaviors (Páll et al., 2002b), suggesting that androgens do not directly inhibit parental care or promote courtship. Instead, it is hypothesized that the drop in androgens after mating may reflect a decreased need for spiggin (Mayer et al., 2004), the proteinaceous glue produced in the kidney that males use for nest construction. Spiggin production is stimulated by androgens (Borg et al., 1993; Jakobsson et al., 1996, 1999). Therefore, a decline in androgens may be related to a decline in spiggin production and nest gluing behavior. We predict that ecotypic differences in gluing behavior will mirror ecotypic differences in androgen levels.

Using quantitative liquid chromatography coupled with mass spectroscopy (LC-MS/MS), we quantified five androgens in males from both ecotypes at three reproductive stages: nesting, immediately after mating (0 days post-fertilization, dpf), and 4 dpf. Sampling at these three stages allowed us to capture the transition from prioritizing mate attraction to parental care that occurs after mating (0 dpf) in commons. At 4 dpf, commons are exclusively focused on parenting and show high levels of fanning behavior (Behrens et al., 2024). Whites were collected at all three stages to compare with commons, although male whites remained ready to mate across stages and never provided parental care. In addition to androgens, we also measured estrogens, progestogens, and corticosteroids, and report those results here. Progestogens are associated with male sexual behavior (Witt et al., 1994; Andersen and Tufik, 2006; Ahamed et al., 2024), so ecotypic differences in progestogens could reflect how whites prioritize mating across stages, whereas commons switch from mating to parenting. Glucocorticoids, which influence energy mobilization and stress (reviewed in Milla et al., 2009), are important in regulating how parents manage the increased energy demands of care (Alonso-Alvarez and Velando, 2012), and how they manage stressors – parents may be more resilient to stressors to some degree, but high levels of stress can lead to decreased care or offspring abandonment (Kaplan et al., 2025). Ecotypic divergence in glucocorticoid levels could tell us if and how the stress response axis may be associated with the loss of care. Our results prompt hypotheses about what endocrine building blocks were tweaked during evolution to construct whites’ and commons’ divergent reproductive strategies.

## 2. METHODS

### 2.1. Animals

We collected adult threespine stickleback fish (*Gasterosteus aculeatus*) of the caregiving common ecotype (“commons”) and non-caregiving white ecotype (“whites”) from two wild populations in Canada in the summers of 2018-2021. Commons were collected from Cherry Burton Road, New Brunswick, Canada (N 46° 01.516’, W 64° 06.150’), and whites were collected from Canal Lake, Nova Scotia, Canada (N 44° 29’54.0”, W 63° 54’09.1”). Collections were approved by the Canadian Department of Fisheries and Oceans Maritime under permit #343930 held by Drs. Anne Dalziel and Laura Weir at Saint Mary’s University. These ecotypes are likely experiencing low hybridization rates at sites where they occur in sympatry (Blouw, 1996; Samuk, 2016). Therefore, these two collection sites where ecotypes generally do not occur in sympatry were selected to reduce the chances of accidentally collecting hybrids. Embryos were generated at Saint Mary’s University and shipped to the University of Illinois Urbana-Champaign, where fish were raised for at least one generation under common garden conditions in the lab before beginning experiments.

Fish were reared in single-family, mixed-sex tanks outfitted with gravel and artificial plants. Experiments were conducted over two summers, from May to July 2021 and June to July 2022, using fish that were sexually mature and approximately one year old. During experiments, fish were maintained in brackish water in light and temperature conditions that simulate summer (10 ppt, 20°C, 16 h light: 8hr dark photoperiod). Fish were fed a mixture of frozen bloodworms, mysis shrimp, and cyclops once a day before beginning experiments, at approximately 7:00 AM. All procedures were approved by the Institutional Animal Care and Use Committee (Protocol #18080 in 2021 and #21031 in 2022).

### 2.2. Behavior

At the beginning of the summers of 2021 and 2022, we visually identified males in breeding condition by their coloration: red throats and blue eyes in commons, and white body coloration in whites. Males had never mated before. Selected males were moved from group housing to individual tanks (36L x 33W x 24H cm) outfitted with gravel, a fake plant, and nesting materials (filamentous *Cladophora* algae and a box of sand). Each male was randomly assigned to one of three stages: nesting, 0 days post-fertilization (0 dpf), or 4 days post-fertilization (4 dpf). Across the two years, we collected data on a total of 22 males per ecotype per stage, except nesting commons, for which we collected 23 males (133 males overall). Once males were relocated to their new tanks, they were left undisturbed overnight to acclimate to their new tanks and start constructing a nest before beginning behavioral observations the next morning.

The following day, we began courtship trials following the methods of Behrens et al., (2024). Briefly, each male was presented with a gravid female of the same ecotype approximately once per day to check for the presence of a completed nest (for males assigned to the nesting stage) or to encourage males to mate (for males assigned to the 0 dpf and 4 dpf stages). Nesting behavior was quantified in 2021 but not in 2022 due to time constraints. Because behavior data is missing for over half our nesting males, here we focus only on the behavior of males collected at 0 and 4 dpf. A detailed account of ecotypic differences in nesting and other behaviors at these stages is reported in Behrens et al., (2024).

We observed and quantified the behavior of males assigned to the 0 and 4 dpf stages to (1) ensure males were at the correct stage to be euthanized for hormone collection (i.e., had successfully mated at 0 dpf, and were actively caring for embryos at 4 dpf (commons only)) and (2) explore the proposed relationship between gluing behavior and 11-KT (Mayer et al., 2004; Páll et al., 2005). Males assigned to 0 dpf or 4 dpf were exposed to gravid females daily until they successfully mated. When a mating occurred, the female was immediately removed, and males were observed for 15 minutes. We quantified behavior using Jwatcher (Blumstein et al., 2006). Here, we focus on gluing behavior because this behavior may be associated with the decline in androgens in caring stickleback (Mayer et al., 2004; Páll et al., 2005). During gluing, males glide over the nest, tail raised, and deposit threads of a sticky protein (spiggin) to fasten nesting material (van Iersel, 1953). We quantified gluing by counting the number of times males performed the behavior during the 15-minute observation period. Fanning, the main measure of parenting in stickleback (van Iersel, 1953), also differs between ecotypes (Behrens et al., 2024), but this behavior is not sensitive to changes in androgens (Páll et al., 2002b). Therefore, we do not present data on fanning in the current study. We continued to observe males assigned to the 4 dpf stage for another four days after they had mated, once a day for 15 minutes, resulting in a total of five observations per male.

All males were euthanized at their assigned stage via rapid decapitation between 9:00 and 13:30 CST to minimize hormonal variations due to time of day. We euthanized nesting males when they had completed their nests but had not mated. A nest was considered complete when males showed nesting behaviors in the presence of a female (i.e., during a courtship trial) (Behrens et al., 2024). We then waited 24 hours after males had seen a female to euthanize them, so they were not influenced by the recent experience with courtship. We sacrificed 0 dpf males 45 minutes after they had successfully courted a female and fertilized eggs. We sacrificed 4 dpf males four days after they had fertilized eggs. For all males, brains were dissected out and set aside for another study and bodies (the entire carcass minus the brain) were immediately flash-frozen on dry ice and stored at −80°C for hormone extraction.

### 2.3. Steroid hormone extraction and quantification

Hormones were extracted from whole bodies (minus the brain) and analyzed via LC-MS/MS. Unlike immunoassays, where cross-reactivity of antibodies to multiple steroids can confound interpretation (Soldin and Soldin, 2009; Ketha et al., 2014), LC-MS/MS can measure steroids of interest with great specificity and allows for the measurement of several steroids simultaneously within a single sample (Soldin and Soldin, 2009; Ketha et al., 2014).

Here, we took advantage of LC-MS/MS to quantify a panel of steroids in whites and commons. Due to the small size of our stickleback (in the lab, adult fish typically weigh about one gram), we chose to quantify steroids in bodies instead of plasma. Steroid levels in whole bodies are correlated with plasma (Graham et al., 2018; Nouri et al., 2020), suggesting this method provides an accurate snapshot of circulating hormone levels when a sufficient volume of plasma cannot be collected. However, it is important to keep in mind that steroids in whole-body extracts reflect not only circulating steroids but also steroids present in tissues, including gonads. While 11-KT is the major androgen in stickleback blood, 11-KA is the major androgen in the testes (Borg and Mayer, 1995).

Hormone extraction and quantification were performed following the methods of (McGhee et al., 2020), except for the homogenization step, which we modified to accommodate our sample type (whole bodies instead of eggs). Extractions were performed in 2023 after samples from both years had been collected. To extract hormones, we coarsely chopped bodies with scissors and ground them into a fine powder using a mortar, pestle, and liquid nitrogen. The weight of each sample was recorded. Steroids were then extracted from each body with methanol. We added 4 mL of methanol to the powdered sample, then vortexed for 60 sec. Samples were stored overnight at −20°C to precipitate neutral lipids and proteins. The following day, samples were spun down (2000 rpm, 4°C) for 20 minutes. The methanol supernatant was collected and diluted with MilliQ water to bring the volume to 50 mL for each sample.

Then, hormones were extracted from the methanol and water solution using solid-phase extraction. C18 Sep-pak cartridges (Waters Cat# WAT020515) were charged with 5 mL of methanol and then rinsed with 5 mL of MilliQ water. Samples were added to the cartridges and pulled through at approximately 2 mL/min with a vacuum. Hormones were eluted with 5 mL of diethyl ether and dried in a fume hood overnight until the ether had evaporated. Dried samples of extracted hormones were stored at −80°C until analysis.

Samples were analyzed using LC-MS/MS by the Carver Metabolomics Core of the Roy J. Carver Biotechnology Center, University of Illinois Urbana-Champaign, with a targeted steroids assay, as previously described (McQueen et al., 2024). We tested for a panel of 35 steroids (see Supplementary Tables 1 and 2 for a complete list of the steroids on this panel). Only free steroids, i.e., unbound to carrier proteins and thought to be biologically active, were quantified. Before extraction, samples were spiked with labeled surrogate internal standards. Following extraction, a 10 μL injection was executed on an Agilent 1290 Infinity II UHPLC system (Agilent, Santa Clara, CA, USA), with a Waters Acquity C18 BEH 2.1 x 150mm, 1.7um; Column Temp - 60°C; flow rate 400 μL/min. Mobile phases consisted of 0.1% formic acid in water (A) and 0.1% formic acid in acetonitrile (B). Data was collected on a Sciex 6500+ triple quadrupole mass spectrometer (Sciex, Framingham, MA, USA). Data was acquired in positive mode using Sciex Analyst v1.7.3. Peak integration and quantitation using calibration curves adjusted for internal standards were done using MultiQuant v3.1 software. Steroid chemical standards, used to create calibration curves for quantification and as internal standards, were acquired from Cayman Chemical (Ann Arbor, MI, USA) and Sigma Aldrich (St. Louis, MO, USA).

### 2.4. Data analysis

Statistical analyses and data visualization were conducted using R Version 4.4.2 (R Core Team, 2024). Steroids were converted to concentrations (ng of steroid per gram of fish) based on the mass of each fish recovered after grinding it into a powder for extraction. We focused our analyses on steroids that were detected in >50% of males per ecotype per stage per year. Of the 35 steroids in our panel, 11 met this criterion. This included five androgens (androstenedione (A4), 11-OH-androstenedione (11-OHA4), testosterone (T), 11-ketoandrostenedione (11-KA), 11-ketotestosterone (11-KT)), one estrogen metabolite (2-methoxyestradiol (2-ME2)), two glucocorticoids (cortisol, cortisone), one mineralocorticoid (aldosterone), and two progesterone metabolites (11β-OH-progesterone (11β-OHP), pregnanolone). Sample sizes for each steroid for each year, ecotype, and stage are provided in Supplementary Tables 1 and 2. One outlier, a white collected at 4 dpf in 2021, was excluded from the analysis and is excluded from the supplementary tables. When an individual had missing data for a steroid (refer to Supplementary Tables 1-2 and figure captions for sample sizes), the missing value was treated as “NA”; we did not perform missing value imputation. For the 11 steroids that met the criteria, one common was missing a value for 11-OHA4, two whites were missing values for pregnanolone, and three whites and one common were missing values for 11β-OHP. Missing data typically occurred for hormones with low median values, suggesting that these missing values represent hormone levels below the detection threshold for LC-MS/MS.

First, we asked how these 11 steroids varied across reproductive stages and between ecotypes. All hormones were first log-transformed, except 11-KT, A4, and 11β-OHP, which were square-root transformed, so residuals would better fit a normal distribution. Note that although steroids were transformed for analysis, all figures present the untransformed values to allow for easier interpretation and visual comparison among steroids. We ran separate linear mixed models for each hormone, including ecotype, stage, and their interaction as fixed effects and year as a random effect using the *lmer* function in the *lme4* package (Bates et al., 2015). When the interaction between ecotype and stage was non-significant, it was dropped from the model. We tested for the significance of fixed effects using the *anova* function in the *lmerTest* package (Kuznetsova et al., 2017) with the Satterthwaite method to estimate degrees of freedom. P-values from these final models were corrected using the Bonferonni-Holm method to account for the fact that we ran multiple separate models on steroids measured in the same individuals. All model p-values remained significant after correction, so we proceeded without correcting the p-values from linear mixed models (i.e., p-values reported in Tables 1-3). We investigated significant effects with post hoc pairwise comparisons using estimated marginal means (EMM) with the *emmeans* function in the *emmeans* package (Lenth, 2024). The resulting p-values from post hoc tests were adjusted for multiple testing using the Tukey method. Effect sizes (Cohen’s d) were calculated for each pairwise comparison using the *eff_size* function in the *emmeans* package. Post hoc test results and effect sizes are reported in Supplementary Tables 3-5.

**Table 1.** Results from linear mixed models testing for effects of ecotype and reproductive stage on the levels of five androgens and one estrogen. Hormone levels were transformed (A4 and 11-KT square-root others log-transformed) before analysis to better fit the assumptions of normality, and models included a random effect of year because data were collected over two years. P-values from all models survived correction; here, we present uncorrected p-values. Significant p-values (p<0.05) are bolded. Degrees of freedom (numerator, denominator) were estimated using the Satterthwaite method. Post hoc test results are reported in Supplementary Table 3.

| Steroid | *F* | df | P |
| --- | --- | --- | --- |
| **Androstenedione (A4)** |  |  |  |
| *ecotype* | 29.041 | 1, 125.02 | **<0.001** |
| *stage* | 3.530 | 2, 125.01 | **0.032** |
| *ecotype\*stage* | 6.967 | 2, 125.00 | **0.001** |
| **11-OH-Androstenedione (11-OHA4)** |  |  |  |
| *ecotype* | 64.870 | 1, 124.00 | **<0.001** |
| *stage* | 10.175 | 2, 124.00 | **<0.001** |
| *ecotype\*stage* | 17.151 | 2, 124.00 | **<0.001** |
| **11-Ketoandrostenedione (11-KA)** |  |  |  |
| *ecotype* | 0.388 | 1, 128.00 | 0.534 |
| *stage* | 4.579 | 2, 128.00 | **0.012** |
| **11-Ketotestosterone (11-KT)** |  |  |  |
| *ecotype* | 33.201 | 1, 125.00 | **<0.001** |
| *stage* | 5.576 | 2, 125.00 | **0.005** |
| *ecotype\*stage* | 9.321 | 2, 125.00 | **<0.001** |
| **Testosterone (T)** |  |  |  |
| *ecotype* | 26.068 | 1, 125.01 | **<0.001** |
| *stage* | 7.450 | 2, 125.01 | **0.001** |
| *ecotype\*stage* | 6.792 | 2, 125.00 | **0.002** |
| **2-Methoxyestradiol (2-ME2)** |  |  |  |
| *ecotype* | 58.453 | 1, 125 | **<0.001** |
| *stage* | 9.075 | 2, 125 | **<0.001** |
| *ecotype\*stage* | 15.882 | 2, 125 | **<0.001** |

This univariate approach, running separate models on each hormone, allowed us to investigate how levels of each hormone differed across stages and ecotypes. We also conducted exploratory analyses using Principal Components Analysis, but by combining data from different hormones together, PCA made it difficult to interpret ecotype and stage effects on specific hormones of interest (such as 11-KT and 11-KA). MANOVA was inappropriate for our data, as the number of response variables exceeded the sample size per group when year, ecotype, and stage were included.

It has been suggested that the effects of progestogens on behavior, i.e., whether they inhibit the actions of androgens or act synergistically to promote male sexual behavior, depend on the relative levels of progestogens to androgens (Wagner, 2006). Therefore, we took advantage of the fact that both androgens and progestogens were measured in the same individuals in our study and calculated the ratio of progestogens to androgen for each individual male. Specifically, we divided the levels of each progestogen metabolite (11β-OHP and pregnanolone) by 11-KT, the major androgen in stickleback (Mayer et al., 2004). Data were log-transformed to better fit a normal distribution, and then linear mixed models were used to test for effects of ecotype and stage, followed by post hoc tests as described above. Because we view this as an exploratory analysis, we only briefly mention the results here; the rest is provided in the Supplement (Supplementary Figure 1, Supplementary Tables 6 and 7).

Next, we leveraged behavioral data from the males euthanized at 4 dpf (n=22 per ecotype) to test the hypothesis that 11-KT levels are associated with levels of gluing behavior, as proposed previously (Mayer et al., 2004; Páll et al., 2005). First, we explored how gluing behavior changed over time in whites and commons, predicting that if gluing behavior followed 11-KT levels, gluing would decline over time in commons but remain steady in whites. To test this, we ran a negative binomial generalized linear mixed model testing for effects of ecotype, days post-fertilization (dpf), and their interaction on the number of times a male glued from 0 to 4 dpf. We included a random effect of year and a random effect of male ID to account for repeated measures. Gluing behavior was measured five times in each male, from 0 dpf, immediately after mating, until 4 dpf, the day males were euthanized. Significant effects were investigated with post hoc tests as described above. Post hoc test results and effect sizes are reported in Supplementary Table 8.

Finally, we examined relationships between gluing and 11-KT at the individual level by testing for correlations between these two variables using the *cor.test* function in the base *stats* package with the Pearson method. We focus on gluing behavior in males collected at 0 and 4 dpf. Because gluing behavior changes over time in commons and differs between ecotypes (see above), we ran separate correlations for each ecotype and stage.

## 3. RESULTS

### 3.1. Levels of androgens (except 11-KA) dropped as commons began providing care, but remained high in whites

We predicted that levels of androgens would decline after mating in the caregiving common ecotype and remain high in the non-caregiving white ecotype. Consistent with this prediction, levels of four androgens (T, 11-KT, A4, 11-OHA4) and one estrogen metabolite (2-ME2) declined across reproductive stages after fertilization in commons, but not whites (Table 1, Fig. 1). In commons, levels of these hormones were significantly higher when males were nesting than when parenting (nesting vs. 4 dpf: T, 11-KT, 11-OHA4, 2-ME2, p<0.001; A4, p=0.001). Additionally, levels of T were greater in nesting commons than post-mating commons that were just beginning to provide care (nesting vs. 0 dpf: p=0.042), and levels of A4, 11-OHA4, and 2-ME2 were greater in commons immediately after mating than when parenting (0 dpf vs. 4 dpf: A4, p=0.029; 11-OHA4 and 2-ME2, p<0.001).

**Figure 1.**
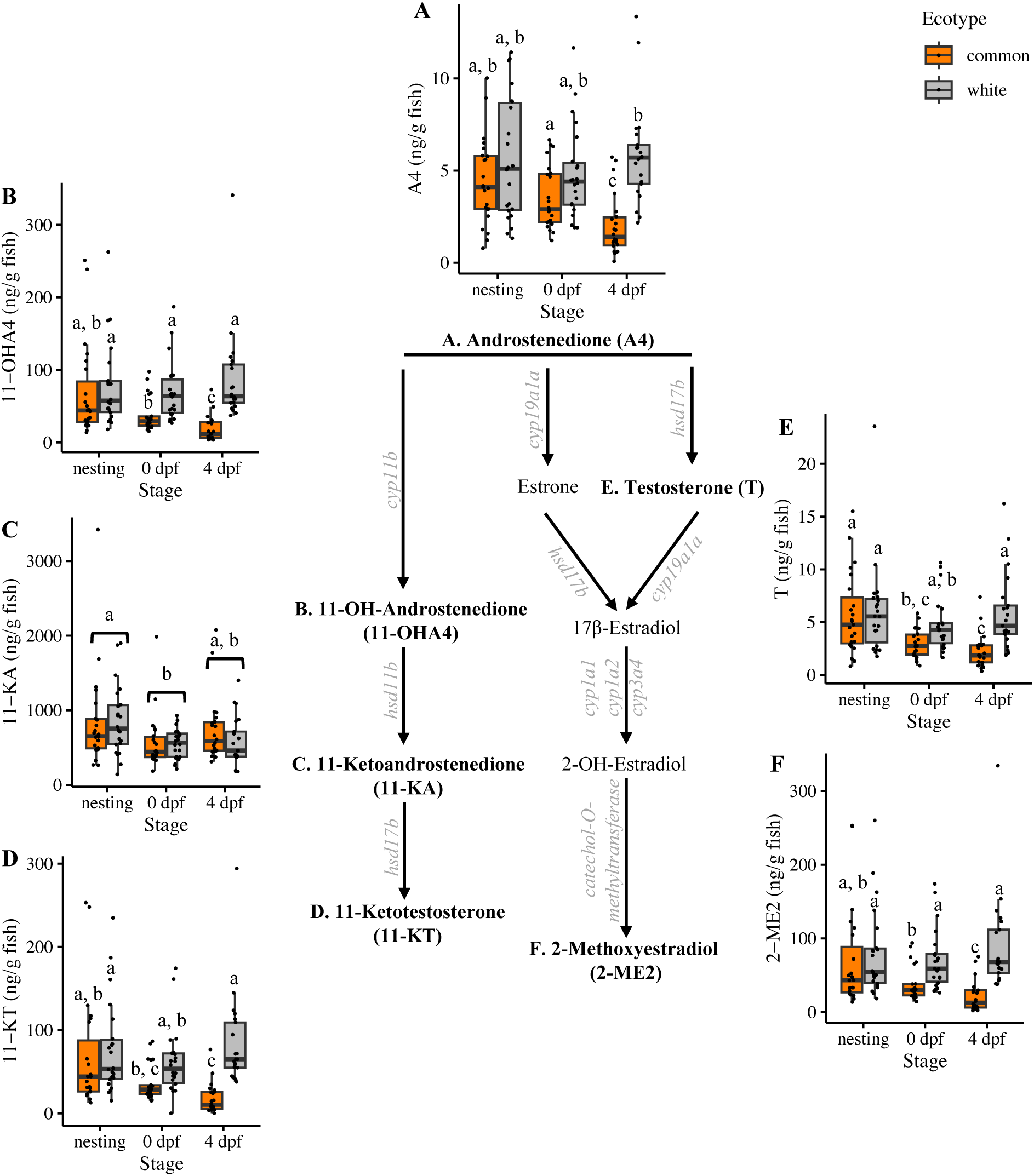
Levels of four androgens and one estrogen metabolite declined in commons after mating; these steroids remained high across all stages in whites. 11-KA was detected at much higher levels than other androgens (note the scale on the Y axis) and showed a different pattern across stages. Levels of each hormone (A-F) are plotted beside a proposed steroidogenic pathway for teleosts, focusing on the six steroids of interest. A4 → 11-OHA4 → 11-KA → 11-KT is the primary steroidogenic pathway in stickleback testes, although the final step (11-KA → 11-KT) primarily occurs in blood (Mayer et al., 1990). Androstenedione can be metabolized into many other steroids in stickleback, including T (Borg et al., 1992). One alternative pathway in teleosts (A4 conversion to estrone and T) is shown here (Tenugu et al., 2021). To our knowledge, the biosynthesis of 2-ME2 from 17β-estradiol is not established in fishes. Therefore, here we show the proposed pathway for mammals (Kumar et al., 2016). Steroids measured in our study are shown in bold font in the pathway; steroidogenic enzymes are in gray italic font. Sample sizes for all hormones except 11-OHA4 were the same: n=23 nesting commons, n=22 nesting whites, n=22 0 dpf commons, n=22 0 dpf whites, n=22 4 dpf commons, and n=21 4 dpf whites. For 11-OHA4, due to missing data for one male, n=21 for 4 dpf commons. For this and subsequent graphs, boxplots show the median (thick black bar), the first and third quartiles (bottom and top edges of box), and 1.5 times the interquartile range (whiskers). Commons are shown in orange, and whites are shown in grey. Each point represents an individual male. Different letters denote significant differences (p<0.05) between groups in estimated marginal means (EMM) post hoc tests after Tukey correction.

In contrast, levels of these five hormones (T, 11-KT, A4, 11-OHA4, and 2-ME2) remained elevated across stages in whites, resembling the levels detected in nesting commons (Table 1, Figure 1). Whites had higher levels of 11-OHA4 and 2-ME2 than commons after mating (0 dpf common vs. white: 11-OHA4, p=0.018; 2-ME2, p=0.029) and higher levels of all five hormones at 4 dpf than commons (4 dpf common vs. white: p<0.001 for all).

Levels of 11-KA were approximately 10x higher than levels of the other detected androgens (e.g., the median level of 11-KT detected across all males was 43.88 ± 4.53, and 11-KA was 592.91 ± 39.12 (median ± standard error ng/g)). 11-KA showed a different pattern across stages and ecotypes than the other androgens (Table 1, Fig. 1). 11-KA levels were higher at the nesting stage than at 0 dpf in both ecotypes (p=0.009), possibly reflecting the important role of androgens in the initiation of reproduction (Mayer et al., 2004). While androgens upstream and downstream of 11-KA showed a decline in commons after 0 dpf, levels of 11-KA did not differ between ecotypes. This difference between 11-KA and the other androgens may be due, in part, to tissue differences in androgen levels. 11-KA is present at low levels in blood but is the most prevalent androgen in testes, whereas the opposite is true for 11-KT (Borg et al., 1989; Mayer et al., 1990). It is possible that the between-stage differences in 11-KA reflect patterns of steroid regulation in testes, and the patterns for 11-KT, the other androgens, and one estrogen reflect patterns in blood. Such tissue-specific regulation of androgens has been described recently in ruffs (Loveland et al., 2025) and has important consequences for male reproductive behavior.

### 3.2. A link between 11-KT and gluing behavior

11-KT is known to promote spiggin (glue) production in stickleback (Borg et al., 1993; Jakobsson et al., 1996, 1999). Therefore, we expected a decline in androgens over time to be associated with a decline in gluing behavior. To explore this, we leveraged behavioral data on individual males observed once daily from 0 dpf when they mated to 4 dpf when they were euthanized.

We detected a significant interaction between ecotype and stage (dpf) on levels of gluing behavior (Fig. 2A: Type III Wald Chi-square= 16.370, df=4, p=0.003); gluing behavior declined after mating in commons but not whites. Common males glued less at 4 dpf than 0 dpf (p<0.0001), but levels of gluing behavior in male whites did not significantly change over time (0 dpf vs. 4 dpf: p=0.979). Also consistent with the ecotypic differences in 11-KT, commons tended to glue less than whites at 4 dpf (commons vs. whites: p=0.085).

**Figure 2.**
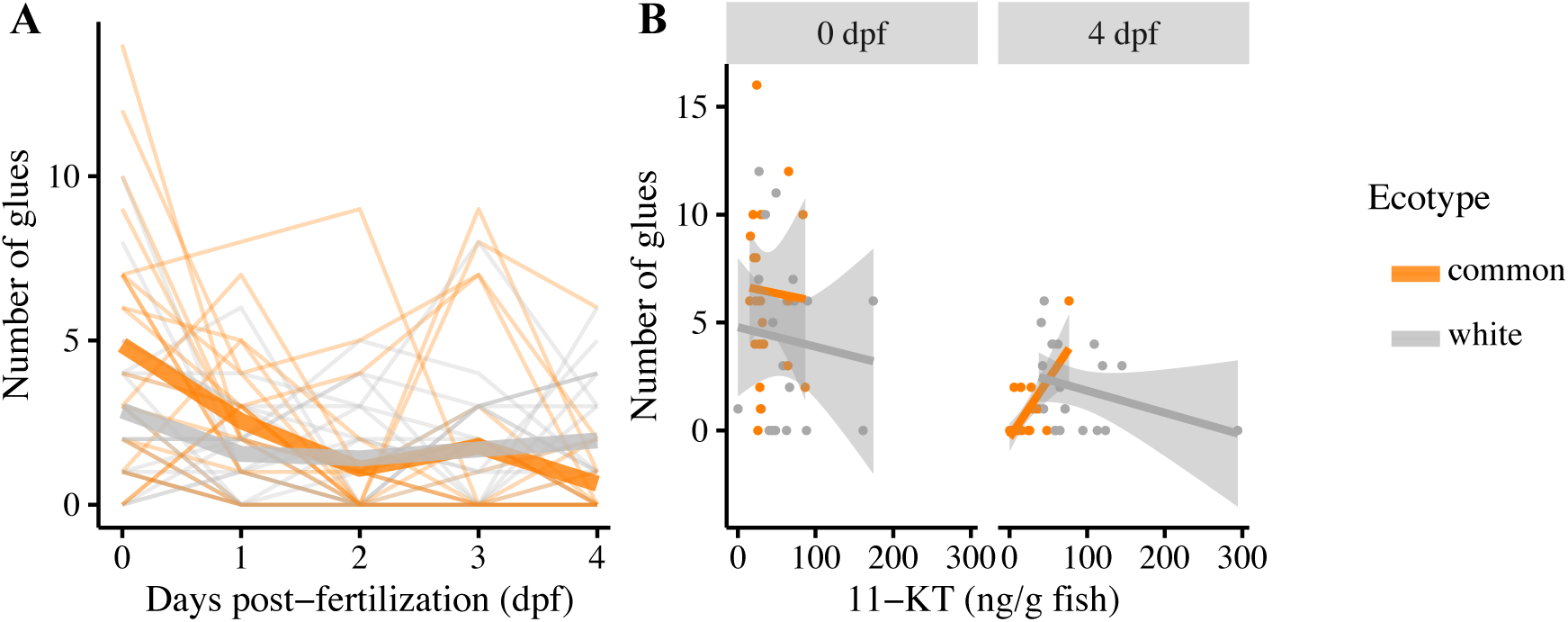
In the common ecotype, gluing behavior declined over time as 11-KT declined, and gluing behavior was positively correlated with 11-KT at 4 dpf. *(A)* Data show the number of glues performed by each male during a once-daily 15-minute observation. Gluing behavior declined over time in commons but not in whites. Each thin line represents the gluing behavior of a single male tracked from the day they mated (0 dpf) until 4 dpf when males were euthanized and collected for hormone quantification. Thick lines represent ecotype averages over time. Only behavior of males collected at 4 dpf are shown, as the others were sacrificed at or before 0 dpf. *(B)* Commons with higher 11-KT at 4 dpf glued their nests more; no significant relationship was detected in whites at 0 or 4 dpf, or commons at 0 dpf. Each point represents one male, and the gray shadow represents 95% confidence intervals.

We examined patterns at the individual level to further explore a possible link between levels of 11-KT and gluing behavior. We tested for correlations between gluing and 11-KT levels in males of each ecotype at 0 dpf and 4 dpf. Commons with higher 11-KT glued more at 4 dpf, but not at 0 dpf (Fig 2B: 0 dpf, r= −0.041, p=0.857, n=22; 4 dpf, r= 0.698, p=0.0003, n=22), providing some support that 11-KT promotes spiggin production and gluing in commons. We did not detect a relationship between 11-KT and gluing in whites (Fig. 2B: 0 dpf, r=-0.096, p=0.670, n=22; 4 dpf, r=-0.300, p=0.187, n=21).

### 3.3. Levels of progestogens were generally lower in whites and after mating (0 dpf)

Progestogens may promote sexual behavior (Witt et al., 1994; Andersen and Tufik, 2006; Ahamed et al., 2024). Therefore, we predicted progestogen levels would differ between male whites and commons, which prioritize mating differently across stages. The two progesterone metabolites we detected, pregnanolone and 11β-OHP, showed similar but not identical patterns across stages and ecotypes (Table 2, Fig. 3). Pregnanolone levels were lower in whites than commons (Table 2, main effect of ecotype, Fig.3B inset). 11β-OHP levels were also lower in whites than commons (Fig. 3A), but this effect was detected only at the nesting stage and at 4 dpf (nesting common vs. white: p=0.005; 4 dpf common vs. white: p=0.012). Across stages, pregnanolone levels were lower at 0 dpf than the other two stages (0 dpf vs nesting and 0 dpf vs 4 dpf: p<0.001). Levels of 11β-OHP were also lower at 0 dpf than the other two stages, but only in the common ecotype (commons 0 dpf vs. nesting and 0 dpf vs. 4 dpf: p<0.001). Levels of 11β-OHP did not significantly differ across stages in whites.

**Figure 3.**
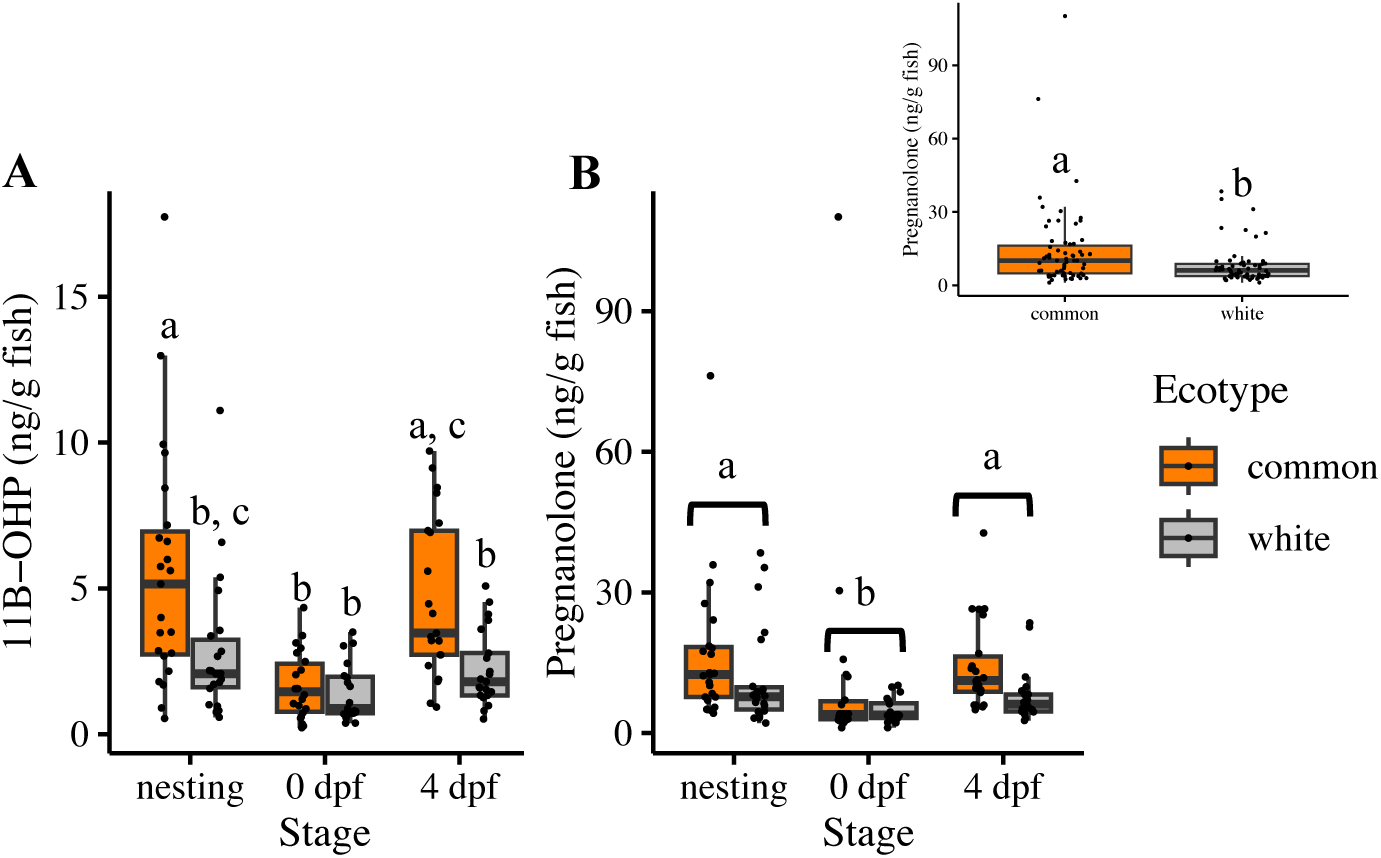
Progestogens differed between ecotypes and across stages. *(A)* 11β-OH-Progesterone (11β - OHP) showed a significant interaction effect between stage and ecotype. *(B)* Pregnanolone showed a main effect of stage (main plot) and ecotype (inset). Different letters denote significant differences between groups. Due to missing data, sample sizes for these two hormones differed slightly. For 11β - OHP n=23 nesting commons, n=22 nesting whites, n=22 0 dpf commons, n=19 0 dpf whites, n=21 4 dpf commons, and n=21 4 dpf whites. Samples sizes were the same for pregnanolone except for 0 dpf whites (n=20) and 4 dpf commons (n=22).

**Table 2.** Results from linear mixed models testing for effects of ecotype and reproductive stage on the levels of two progestogens. Hormone levels were transformed (11β-OHP was square-root and pregnanolone was log-transformed) before analysis to better fit the assumptions of normality, and models included a random effect of year because data were collected over two years. P-values from all models survived correction; here, we present uncorrected p-values. Significant p-values (p<0.05) are bolded. Degrees of freedom (numerator, denominator) were estimated using the Satterthwaite method. Post hoc test results are reported in Supplementary Table 4.

| **Steroid** | **F** | **df** | **P** |
| --- | --- | --- | --- |
| **11β-OH-Progesterone (11β-OHP)** |  |  |  |
| *ecotype* | 17.984 | 1, 121.00 | **<0.001** |
| *stage* | 17.564 | 2, 121.07 | **<0.001** |
| *ecotype\*stage* | 3.108 | 2, 121.02 | **0.048** |
| **Pregnanolone** |  |  |  |
| *ecotype* | 12.043 | 1, 125.00 | **0.001** |
| *stage* | 16.560 | 2, 125.01 | **<0.001** |

We also explored progestogen-to-androgen ratios (levels of 11β-OHP and pregnanolone relative to 11-KT) because the effects of progestogens on male sexual behavior may depend on androgen levels. Progestogen to androgen ratios were low when levels of male sexual behavior were high, i.e., across all stages in whites and at the nesting and 0 dpf stages in commons (Supplementary Fig. 1, Supplementary Table 6). In contrast, progesterone to androgen ratios were relatively high (approximately 7x) when levels of sexual behavior were low, i.e. in parenting commons, which had low levels of 11-KT and high levels of progestogens (Supplementary Fig. 1, Supplementary Table 7).

### 3.4. Corticosteroids were elevated after mating (0 dpf) in both ecotypes

Parenthood is associated with changes in energy homeostasis and stress response, which are regulated by glucocorticoids (Milla et al., 2009). Therefore, we predicted that commons may show elevated glucocorticoids relative to whites at 0 and/or 4 dpf, if corticosteroids are involved in mobilizing energy for parenting. Alternatively, whites may show elevated glucocorticoids relative to commons at 0 dpf if offspring abandonment is triggered by a stress response in whites. We found no evidence to support either prediction.

We detected two glucocorticoids (cortisol and cortisone) and one mineralocorticoid (aldosterone). Unlike the androgens and progestogens, we did not detect a significant interaction effect of ecotype and stage on any corticosteroids, but there was a significant main effect of stage (Table 3). Levels of all three corticosteroids were significantly elevated in both ecotypes after mating compared to 4 dpf (Fig. 4, 0 dpf vs. 4 dpf: cortisol, p<0.001; cortisone, p=0.007; aldosterone, p=0.011). For cortisol only, levels were also significantly higher at 0 dpf than at the nesting stage (p=0.019).

**Table 3.**
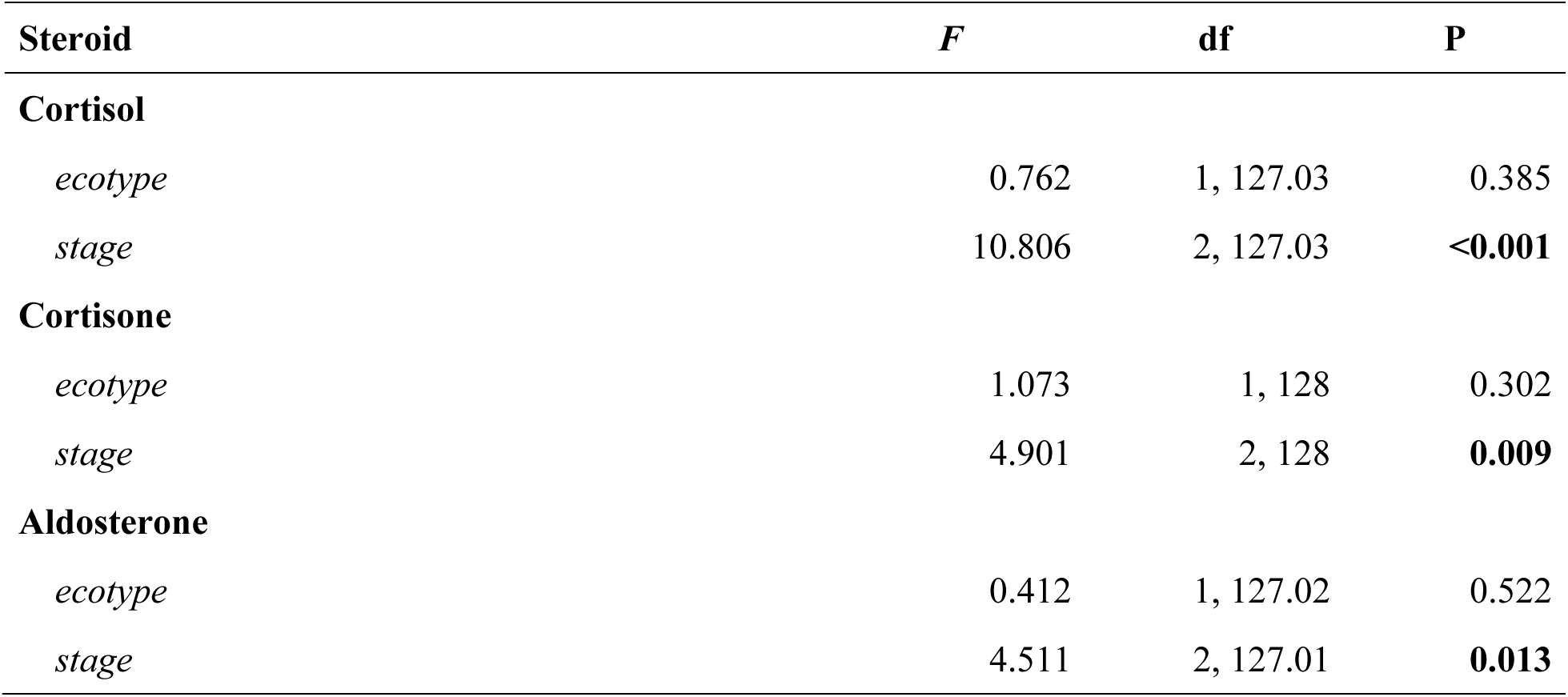
Results from linear mixed models testing for effects of ecotype and reproductive stage on the levels of three corticosteroids. Hormone levels were log-transformed before analysis to better fit the assumptions of normality, and models included a random effect of year because data were collected over two years. P-values from all models survived correction; here, we present uncorrected p-values. Significant p-values (p<0.05) are bolded. Degrees of freedom (numerator, denominator) were estimated using the Satterthwaite method. Post hoc test results are reported in Supplementary Table 5.

**Figure 4.**
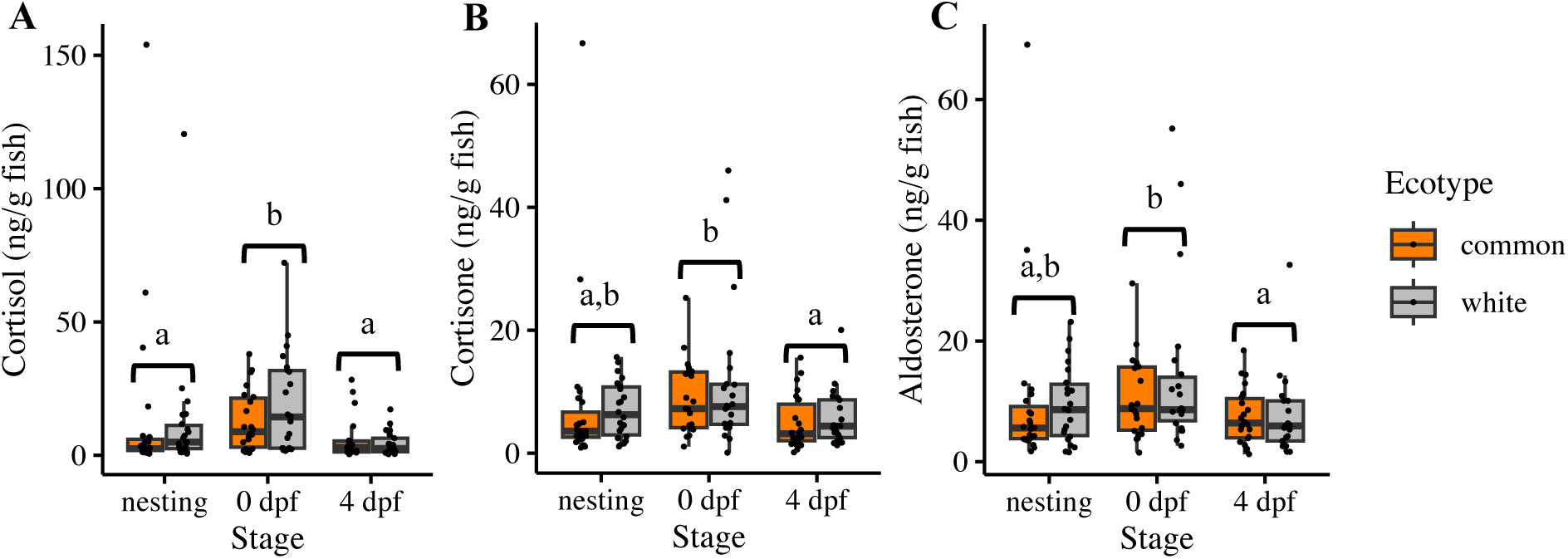
Corticosteroid levels were higher after mating in both ecotypes. Patterns were similar across all three corticosteroids detected: *(A)* cortisol, *(B)* cortisone, and *(C)* aldosterone. Levels of corticosteroids did not significantly differ between ecotypes at any stage; different letters denote significant differences between stages pooled across ecotypes. Sample sizes for the three hormones were the same: n=23 nesting commons, n=22 nesting whites, n=22 0 dpf commons, n=22 0 dpf whites, n=22 4 dpf commons, and n=21 4 dpf whites.

## 4. DISCUSSION

### 4.1. Evidence for ecotypic differences in androgen regulation

Androgens play an important role in male reproductive behavior and trade-offs between parental care and mating (Wingfield et al., 1990; Oliveira et al., 2002). To understand how androgens and other steroids contribute to the evolution of different reproductive strategies, we can compare hormone levels in closely related, behaviorally divergent populations. Here, we take advantage of a pair of recently diverged threespine stickleback ecotypes to gain insights into how endocrine building blocks, particularly those involved in androgen signaling, may have been tweaked in the evolutionary loss of parental care. We predicted that males of the white ecotype that show high rates of courtship behavior (Haley et al., 2019; Behrens et al., 2024) and abandon their offspring soon after fertilization (Blouw and Hagen, 1990; Blouw, 1996) would maintain consistently high levels of androgens. We predicted that males of the common ecotype that stop courting females a few days after mating to provide care to their offspring (Jamieson et al., 1992) would show a drop in androgens after mating. Consistent with this hypothesis, we found that levels of most, but not all, androgens declined in commons after mating and remained high in whites, possibly due to ecotypic differences in the costs and benefits of androgen-mediated traits.

Consistent with our predictions and previous work in stickleback (Páll et al., 2002a) 11-KT declined in commons as they transitioned into caregiving after mating and remained elevated in whites across stages. This suggests that these ecotypes have diverged in how they regulate androgen levels during reproduction. Except for 11-KA, all other androgens (A4, T, 11-OHA4) and one estrogen metabolite (2-ME2) downstream of A4 followed this pattern, which suggests that at least some of these changes occurred early in the steroidogenic pathway in gonad and/or upstream of the gonad. This pattern prompts at least four non-mutually exclusive evolutionary changes.

First, the gonadotropin pituitary hormones (luteinizing hormone (LH) and follicle-stimulating hormone (FSH)) promote androgen production in the testes (Planas, 1995) and decline in parental stickleback after mating, mirroring the changes in 11-KT (Páll et al., 2005). Therefore, one possibility is that the pituitaries of male whites continue to produce LH or FSH after mating and/or maintain greater LH or FSH receptor binding or expression in testes, resulting in consistently high androgen levels.

Second, another possibility is that the primary site of divergence between whites and commons lies in the brain. There might be differences in androgen receptor affinity, binding, or density in the hypothalamus, for example, that lead to differences in gonadotropin release and, ultimately, androgen levels. Previous studies found that male commons increase activation of preoptic oxytocin neurons after mating, but whites do not (Maciejewski et al., in press), and commons have higher pituitary oxytocin peptide levels (Bell et al., 2025). Oxytocin neurons in the preoptic area of the hypothalamus project to the pituitary and can influence gonadotrophs (Trudeau et al., 2024). Therefore, the differential activity of oxytocin neurons in the preoptic area of the hypothalamus could be driving the observed differences in androgen levels.

Third, it is also possible that whites and commons have diverged at an even higher level of organization, for example, at the level of the sensory systems that detect cues that trigger parental behavior. Previous work in stickleback suggests that changes in androgens after mating may be induced by the experience of mating or cues from eggs, as males that mate more times and have more eggs show more rapid declines in 11-KT levels (Páll et al., 2002a). This suggests that there is a mechanism in commons that responds to external cues following mating that triggers a decline in androgens, and whites fail to show this response. Whether this difference in whites is due to a change in the cues themselves (e.g., ecotypic differences in egg olfactory cues), a lack of sensitivity to cues (e.g., differences in males’ olfactory systems), or differences in the processing of sensory cues remains an interesting and open question.

Fourth, there is recent evidence that steroidogenic enzymes in blood are an underappreciated but potentially potent mechanism that can modulate the effects of gonadal steroids on the brain and thereby be related to behavioral variation. For example, there are alternative mating strategies in ruffs that harbor alleles that differ in the efficiency of the 17-beta hydroxysteroid dehydrogenase 2 (hsd17b2) enzyme, which converts testosterone to androstenedione (Loveland et al., 2025). The behavioral strategies characterized by low testosterone (satellites and faeders) harbor allelic variants of the hsd17b2 enzyme that are more efficient and, thereby, more effective in preventing testosterone from reaching the brain. Interestingly, hydroxysteroid dehydrogenase converts 11-KA to 11-KT in blood in male sticklebacks (Mayer et al., 1990). The finding that 11-KT drops in commons but not whites after mating suggests that commons may have lower hydroxysteroid dehydrogenase activity in blood than whites after mating, and/or genetic variation between the ecotypes at that locus, as in the ruffs. Future work quantifying steroidogenic enzyme expression in blood and testes and comparing allelic variation in whites and commons can shed more light on the origins of ecotypic differences in androgen levels and the high levels and lack of ecotype difference in 11-KA.

Importantly, changes at all these levels of biological organization are not mutually exclusive, and it is possible that several mechanisms may be at play. For example, two ecotypes of stickleback that diverged in courtship behavior (Kitano et al., 2008) and 11-KT levels (Kitano et al., 2011) were found to differ in LH receptor and steroidogenic enzyme expression in testes (Kitano et al., 2011). A similar combination of mechanisms may explain the steroid differences in whites and commons. Interactions between androgens and other steroids should also be considered (for example, progestogens, see section 4.3), as ratios of steroid levels can influence feedback on the HPG axis (Trudeau et al., 2024). Future studies should explore these and other possible modulators of androgen levels to understand how commons reduce their androgen levels after mating, and whites do not, thereby gaining insight into the building blocks of behavioral evolution (Rosvall, 2022).

### 4.2. Are differences in androgen regulation related to the need for nest glue (spiggin)?

Changes in 11-KT levels (or a lack thereof in whites) may be related to ecotypic difference in the need for spiggin (Mayer et al., 2004). After mating, caregiving males show decreased courtship, decreased gluing, and increased parental fanning (Páll et al., 2002a, 2005; Behrens et al., 2024). Male whites maintain courtship and gluing behavior after mating and display no parental fanning (Jamieson et al., 1992; Behrens et al., 2024). These patterns may initially lead us to conclude that high androgens are incompatible with parental care, as is the case for several species of birds (Silverin, 1980; Hegner and Wingfield, 1987; Oring et al., 1989; Ketterson et al., 1992; De Ridder et al., 2000; Stoehr and Hill, 2000). However, androgen manipulation studies have shown that while spiggin production is androgen-mediated (Páll et al., 2005), fanning and courtship are not (Páll et al., 2002b). We found that gluing behavior declined in commons after mating as they transitioned into parenting, and commons with higher 11-KT glued more at 4 dpf, consistent with work in other stickleback (Páll et al., 2005). For the first time, we show that the relationship between gluing behavior and 11-KT levels also holds true in the white ecotype; whites maintained steady levels of gluing behavior and 11-KT across stages. This suggests that differences in androgen levels after mating in whites and commons may be related to the costs of maintaining androgens and androgen-mediated traits, like spiggin or coloration (Borg et al., 1993; Jakobsson et al., 1996, 1999; Kurtz et al., 2007). Male commons don’t require bright nuptial coloration to attract mates or spiggin to build and maintain nests after mating. In fact, parental stickleback dismantle their nests in the days following mating, presumably allowing for more embryo ventilation as they develop (van Iersel, 1953). Androgens can have detrimental effects on immunity (Kurtz et al., 2007), and spiggin is energetically costly to produce (Östlund-Nilsson, 2001); therefore, when androgen-mediated traits are no longer beneficial, males may benefit from reducing androgen production and reallocating resources to care, which is itself costly (Smith and Wootton, 1999). Whites maintain elevated androgens across stages, perhaps because, unlike commons, whites continue to court females throughout the breeding season and need to maintain spiggin production to keep nests for potential mates (Jamieson et al., 1992; Blouw, 1996). The benefits of increased reproductive success conferred by maintaining high 11-KT likely outweigh the costs for male whites (Wingfield et al., 2001).

### 4.3. Low progestogen levels may promote sexual behavior

In addition to androgens, we quantified two metabolites of progesterone (pregnanolone and 11β - OH-progesterone (11β-OHP). Both metabolites showed relatively consistent differences between ecotypes and across stages. Levels were lower in whites and at 0 dpf when males had just mated. This is consistent with a growing literature that implicates progesterone in male reproduction and androgen-dependent sexual behavior (Witt et al., 1994; Andersen and Tufik, 2006). Generally, the effects of progesterone on male sexual behavior appear to be dose-dependent; at low doses, progesterone can facilitate male sexual behavior, and high doses can inhibit it (Wagner, 2006). Consistent with this, the progestogens we detected were lowest in both ecotypes immediately after mating (0 dpf), suggesting that these metabolites or their precursors could promote mating behavior when present at low levels.

Interestingly, the effects of progesterone may depend on the relative levels of androgens. Work in green anoles (Young et al., 1991) and zebrafish (Ahamed et al., 2024) suggests that male sexual behavior is only facilitated by progesterone when progesterone levels are low relative to androgen levels. Our LC-MS/MS panel allowed us to explore this idea because we measured both 11-KT and progestogens in each male. Our data support the idea that a low progestogen-to-androgen ratio facilitates sexual behavior, and a high ratio may inhibit it. Progestogen metabolite levels were low relative to 11-KT levels across stages in whites and at nesting and 0 dpf in commons, times when males are expected to show sexual behavior. In commons at 4 dpf, when sexual behavior is expected to subside and parental behavior dominates (Jamieson et al., 1992; Behrens et al., 2024), progestogens were high relative to 11-KT. Perhaps, ecotypic differences in the balance of progestogens and androgens may promote whites keeping sexual behavior “on” across stages, and commons turning sexual behavior “off” after 0 dpf as they transition exclusively into caregiving. It should be acknowledged that these ratios were calculated using progestogen metabolites, not DHP, the major progestogen in teleosts (Scott et al., 2010), so more work is needed to confirm these patterns hold true for DHP. Simultaneously manipulating 11-KT and progestogen levels in breeding males would be a powerful approach to determine how concerted action by these steroids may influence ecotypic differences in male sexual behavior.

### 4.4. No evidence for ecotypic differences in energy homeostasis or stress response

Corticosteroids, unlike androgens and progestogens, showed no differences between ecotypes but did differ across stages. Glucocorticoids mediate energy homeostasis and stress response in fishes (Milla et al., 2009), so the lack of ecotypic differences suggests that the evolutionary loss of care in whites is not associated with changes in energy homeostasis or the stress response.

Parenting is energetically demanding and physiologically costly (Smith and Wootton, 1999; Alonso-Alvarez and Velando, 2012), and in mammals, particularly mothers, the onset of parenting is associated with changes in the stress response and energy homeostasis (Kaplan et al., 2025); evidence in fathers is mixed (Saltzman and Ziegler, 2014). There is also some evidence in birds (Kotrschal et al., 1998; Golet et al., 2004) and fish (Magee et al., 2006) for elevated glucocorticoids in parents relative to non-parents. If high glucocorticoids support energy mobilization to meet the demands of caring, we might expect commons to show higher glucocorticoids than whites at 4 dpf when commons’ parental effort peaks (Behrens et al., 2024). Alternatively, if high glucocorticoids reflect high stress, which can lead to offspring abandonment (Kaplan et al., 2025), we might expect whites to show higher corticosteroids after mating when they spit their offspring out of the nest. Contrary to both predictions, both cortisol and cortisone were elevated at 0 dpf in both ecotypes. This suggests that glucocorticoid levels may respond to a shared experience by whites and commons, such as mating, as shown in mammals (Borg et al., 1991; Villani et al., 2006), and/or courtship. Previous work in stickleback found that cortisol levels were elevated in socially isolated stickleback after brief exposure to a conspecific (Bell et al., 2007). Perhaps the elevated glucocorticoids in our fish were a response to their interaction with a female during courtship, as males were otherwise housed alone.

Also worth noting is that aldosterone, a mineralocorticoid considered absent, or at least not present at any significant level, in fishes (Gilmour, 2005; Milla et al., 2009), was detected in all the males in our study, a testament to the power of LC-MS/MS. Aldosterone has been detected in fishes in at least one other LC-MS/MS study (Nouri et al., 2020). This warrants further investigation to determine what role aldosterone plays in the physiology and behavior of stickleback and fishes more generally.

## 5. CONCLUSIONS

Here, we describe differences in steroid levels across reproductive stages in two behaviorally divergent ecotypes of threespine stickleback as a first step toward understanding how neuroendocrine mechanisms evolve when parental care is lost. Leveraging LC-MS/MS, we quantified a panel of eleven steroids. We demonstrate that this behavioral divergence was not isolated to one key hormone, such as 11-KT. Rather, seven of eleven measured steroids differed between ecotypes, including four androgens, one estrogen metabolite, and two progestogen metabolites. Corticosteroid levels peaked after mating but did not differ between ecotypes, suggesting that in the evolution of reproductive behavior, steroids in the HPG axis may be more likely to be targeted than the hypothalamic-pituitary-adrenal axis (or hypothalamic-pituitary-interrenal (HPI) axis in fishes). However, more work is needed to determine if other aspects of corticosteroid signaling, such as crosstalk between the HPG and HPI (Rosvall et al., 2016), may differ between ecotypes.

We found that androgen levels declined in commons across stages but remained elevated in whites, suggesting that the regulation of steroid levels over time has diverged between ecotypes. Although androgens showed the expected negative relationship with care behavior in commons, androgens are not causally related to care in stickleback (Páll et al., 2002b); therefore, the loss of care in whites is unlikely to be a result of high androgens. But consistently high androgen levels in whites may support the maintenance of sexual behavior as 11-KT does promote nuptial coloration, spiggin production, and possibly gluing behavior (Borg et al., 1993; Jakobsson et al., 1996, 1999; Kurtz et al., 2007). Progestogen levels also remain low in whites across stages, which could, alongside high androgen levels, facilitate the maintenance of high levels of sexual behavior. Future work will explore potential mechanisms for ecotypic differences in androgens, such as ecotypic differences in the HPG axis due to differences in hormones receptors, gonadotropin levels, and/or expression of steroidogenic enzymes like hsd17B. This will ultimately further our understanding of how endocrine building blocks are tweaked by evolution in the early stages of behavioral divergence.

## ACKNOWLEDGEMENTS

We thank Anne Dalziel and Laura Weir for help in the field and members of the Bell Lab for support and helpful feedback. We thank Ryan Paitz, Sarah Winnicki, Sarah Westrick, and Eva Fischer for helpful conversations and guidance on steroid extraction. We thank Annika Bagazinski for her assistance extracting hormones, Tara Pavithran for her role in methods development, Cassie Afseth and Kaithren Garcia for their help with behavioral observations, and the University of Illinois Roy J. Carver Metabolomics Core for LC-MS/MS. During this work, MFM was supported by a Graduate Research Fellowship from the National Science Foundation and an Illinois Distinguished Fellowship from the University of Illinois. The research reported in this publication was supported by an Odum-Kendeigh Summer Research Award from the University of Illinois, an R.C. Lewontin Graduate Research Excellence Grant from the Society for the Study of Evolution, and the NIGMS of the National Institutes of Health [award number 1R35GM139597].

