## Supplementary Materials for "An evolutionary shift to prioritizing mating over care is associated with consistently high androgen levels in male threespine stickleback"

**Supplementary Table 1.** Sample sizes for each of the 35 hormones tested for in our 2021 samples for LC-MS/MS, separated by ecotype and stage. Hormones are sorted by sample size; hormones at the top were detected in all males, and those at the bottom were detected in no males. The eleven hormones that exceeded the minimum sample size threshold (>50% per ecotype per stage) are bolded. These eleven hormones were used to test for the effects of ecotype and stage on hormone levels. 5β-tetrahydrocortisone and β-cortolone could not be differentiated, so they are listed together in one row. Abbreviations: DHP=Dihydroprogesterone, DHEA=dehydroepiandrosterone.

| Hormone | Number of males in which hormone was detected | | | | | |
|---|---|---|---|---|---|---|
| | Common | | | White | | |
| | Nesting | 0 dpf | 4 dpf | Nesting | 0 dpf | 4 dpf |
| **Testosterone** | 11 | 10 | 10 | 10 | 10 | 9 |
| **11-Ketotestosterone** | 11 | 10 | 10 | 10 | 10 | 9 |
| **Androstenedione** | 11 | 10 | 10 | 10 | 10 | 9 |
| **11-OH-Androstenedione** | 11 | 10 | 10 | 10 | 10 | 9 |
| **11-Ketoandrostenedione** | 11 | 10 | 10 | 10 | 10 | 9 |
| **2-Methoxyestradiol** | 11 | 10 | 10 | 10 | 10 | 9 |
| **Cortisol** | 11 | 10 | 10 | 10 | 10 | 9 |
| **Cortisone** | 11 | 10 | 10 | 10 | 10 | 9 |
| **Aldosterone** | 11 | 10 | 10 | 10 | 10 | 9 |
| **Pregnanolone** | 11 | 10 | 10 | 10 | 10 | 9 |
| **11β-OH-Progesterone** | 11 | 10 | 9 | 10 | 9 | 9 |
| 5β-DHP | 11 | 5 | 9 | 6 | 7 | 4 |
| Corticosterone | 8 | 8 | 8 | 5 | 5 | 7 |
| 5β-Dihydrocortisol | 9 | 5 | 3 | 4 | 4 | 2 |
| DHEA | 9 | 3 | 4 | 1 | 0 | 1 |
| Deoxycorticosterone | 4 | 0 | 5 | 1 | 2 | 0 |
| 11-Deoxycortisol | 5 | 1 | 2 | 1 | 1 | 1 |
| 5β-Dihydrocortisone | 2 | 5 | 1 | 1 | 2 | 0 |
| 5β-Tetrahydrocortisone | 1 | 1 | 0 | 0 | 1 | 0 |
| 17α-OH-Progesterone | 0 | 1 | 0 | 1 | 0 | 0 |
| Etiocholanolone | 0 | 1 | 1 | 0 | 0 | 0 |
| Transandrosterone | 0 | 1 | 0 | 0 | 0 | 0 |
| 20β-DHP | 0 | 0 | 0 | 1 | 0 | 0 |
| Estradiol | 0 | 0 | 0 | 0 | 0 | 0 |
| Estrone | 0 | 0 | 0 | 0 | 0 | 0 |
| Progesterone | 0 | 0 | 0 | 0 | 0 | 0 |
| Pregnenolone | 0 | 0 | 0 | 0 | 0 | 0 |
| 20β-Dihydrocorticosterone | 0 | 0 | 0 | 0 | 0 | 0 |
| 11-Tetrahydrocorticosterone | 0 | 0 | 0 | 0 | 0 | 0 |
| 5β-Corticosterone | 0 | 0 | 0 | 0 | 0 | 0 |
| β-Cortol | 0 | 0 | 0 | 0 | 0 | 0 |
| 5β-Tetrahydrocorticosterone | 0 | 0 | 0 | 0 | 0 | 0 |
| 5β-Tetrahydrocortisone/β-Cortolone | 0 | 0 | 0 | 0 | 0 | 0 |
| 5α-DHP | 0 | 0 | 0 | 0 | 0 | 0 |
| 17α-OH-Pregnenalone | 0 | 0 | 0 | 0 | 0 | 0 |

**Supplementary Table 2.** Sample sizes for each of the 35 hormones tested for in our 2022 samples for LC-MS/MS, separated by ecotype and stage. Hormones are sorted by sample size; hormones at the top were detected in all males, and those at the bottom were detected in no males. The eleven hormones that exceeded the minimum sample size threshold (>50% per ecotype per stage) are bolded. These eleven hormones were used to test for the effects of ecotype and stage on hormone levels. 5β-tetrahydrocortisone and β-cortolone could not be differentiated, so they are listed together in one row. Abbreviations: DHP=Dihydroprogesterone, DHEA=dehydroepiandrosterone.

| Hormone | Number of males in which hormone was detected | | | | | |
|---|---|---|---|---|---|---|
| | Common | | | White | | |
| | Nesting | 0 dpf | 4 dpf | Nesting | 0 dpf | 4 dpf |
| **Testosterone** | 12 | 12 | 12 | 12 | 12 | 12 |
| **11-Ketotestosterone** | 12 | 12 | 12 | 12 | 12 | 12 |
| **Androstenedione** | 12 | 12 | 12 | 12 | 12 | 12 |
| **11-Ketoandrostenedione** | 12 | 12 | 12 | 12 | 12 | 12 |
| **2-Methoxyestradiol** | 12 | 12 | 12 | 12 | 12 | 12 |
| **Cortisol** | 12 | 12 | 12 | 12 | 12 | 12 |
| **Cortisone** | 12 | 12 | 12 | 12 | 12 | 12 |
| **Aldosterone** | 12 | 12 | 12 | 12 | 12 | 12 |
| **11-OH-Androstenedione** | 12 | 12 | 11 | 12 | 12 | 12 |
| **11β-OH-Progesterone** | 12 | 12 | 12 | 12 | 10 | 12 |
| **Pregnanolone** | 12 | 12 | 12 | 12 | 10 | 12 |
| 5β-DHP | 12 | 9 | 11 | 10 | 10 | 12 |
| Corticosterone | 6 | 11 | 9 | 7 | 10 | 7 |
| Deoxycorticosterone | 11 | 9 | 7 | 2 | 4 | 4 |
| 5β-Dihydrocortisol | 1 | 9 | 4 | 1 | 7 | 3 |
| 11-Deoxycortisol | 2 | 6 | 3 | 2 | 4 | 3 |
| DHEA | 7 | 2 | 5 | 1 | 2 | 2 |
| 5β-Dihydrocortisone | 1 | 4 | 2 | 1 | 7 | 1 |
| 5β-Tetrahydrocortisone | 1 | 2 | 1 | 1 | 4 | 0 |
| Progesterone | 1 | 0 | 3 | 0 | 1 | 1 |
| 17α-OH-Progesterone | 0 | 1 | 0 | 0 | 2 | 0 |
| Transandrosterone | 0 | 2 | 1 | 0 | 0 | 0 |
| 20β-Dihydrocorticosterone | 0 | 1 | 0 | 0 | 0 | 0 |
| Estradiol | 0 | 0 | 0 | 0 | 0 | 0 |
| Estrone | 0 | 0 | 0 | 0 | 0 | 0 |
| Pregnenolone | 0 | 0 | 0 | 0 | 0 | 0 |
| 11-Tetrahydrocorticosterone | 0 | 0 | 0 | 0 | 0 | 0 |
| 5β-Corticosterone | 0 | 0 | 0 | 0 | 0 | 0 |
| β-Cortol | 0 | 0 | 0 | 0 | 0 | 0 |
| 5β-Tetrahydrocorticosterone | 0 | 0 | 0 | 0 | 0 | 0 |
| 5β-Tetrahydrocortisone/β-Cortolone | 0 | 0 | 0 | 0 | 0 | 0 |
| Etiocholanolone | 0 | 0 | 0 | 0 | 0 | 0 |
| 20β-DHP | 0 | 0 | 0 | 0 | 0 | 0 |
| 5α-DHP | 0 | 0 | 0 | 0 | 0 | 0 |
| 17α-OH-Pregnenalone | 0 | 0 | 0 | 0 | 0 | 0 |

**Supplementary Table 3.** Results of post-hoc tests and effect sizes corresponding to Table 1 (androgens and 2-ME2). Emmeans and effect sizes for pairwise differences in steroid levels are reported on the transformed (sqrt and log) scale. P-values are corrected for multiple testing using the tukey correction and effect size is Cohen's d. Significant p-values ($p<0.05$) are bolded.

| response variable | contrast | estimate | SE | df | t-ratio | p-value | effect size |
|---|---|---|---|---|---|---|---|
| sqrt(A4) | common nesting – white nesting | -0.236 | 0.168 | 125 | -1.403 | 0.7252 | -0.418 |
| | common 0 dpf – white 0 dpf | -0.303 | 0.170 | 125 | -1.781 | 0.4815 | -0.537 |
| | common 4 dpf – white 4 dpf | -1.049 | 0.172 | 125 | -0.6096 | **<0.0001** | -1.860 |
| | common nesting – common 0 dpf | 0.200 | 0.168 | 125 | 1.188 | 0.8421 | 0.354 |
| | common nesting – common 4 dpf | 0.726 | 0.168 | 125 | 4.316 | **0.0005** | 1.287 |
| | common 0 dpf – common 4 dpf | 0.526 | 0.170 | 125 | 3.095 | **0.0287** | 0.933 |
| | white nesting – white 0 dpf | 0.133 | 0.170 | 125 | 0.781 | 0.9702 | 0.236 |
| | white nesting – white 4 dpf | -0.087 | 0.172 | 125 | -0.506 | 0.9959 | -0.154 |
| | white 0 dpf – white 4 dpf | -0.220 | 0.172 | 125 | -1.278 | 0.7966 | -0.390 |
| log(11-OHA4) | common nesting – white nesting | -0.2543 | 0.183 | 124 | -1.388 | 0.7341 | -0.4140 |
| | common 0 dpf – white 0 dpf | -0.6029 | 0.185 | 124 | -3.256 | **0.0179** | -0.9816 |
| | common 4 dpf – white 4 dpf | -1.7381 | 0.190 | 124 | -9.166 | **<0.0001** | -2.8298 |
| | common nesting – common 0 dpf | 0.4005 | 0.183 | 124 | 2.186 | 0.2516 | 0.6521 |
| | common nesting – common 4 dpf | 1.3325 | 0.185 | 124 | 7.188 | **<0.0001** | 2.1695 |
| | common 0 dpf – common 4 dpf | 0.9320 | 0.187 | 124 | 4.973 | **<0.0001** | 1.5174 |
| | white nesting – white 0 dpf | 0.0519 | 0.185 | 124 | 0.280 | 0.9998 | 0.0844 |
| | white nesting – white 4 dpf | -0.1513 | 0.187 | 124 | -0.807 | 0.9657 | -0.2463 |
| | white 0 dpf – white 4 dpf | -0.2031 | 0.187 | 124 | -1.084 | 0.8869 | -0.3307 |
| log(11-KA) | nesting – 0 dpf | 0.339 | 0.113 | 127 | 2.992 | **0.0093** | 0.634 |
| | nesting – 4 dpf | 0.213 | 0.114 | 127 | 1.864 | 0.1536 | 0.398 |
| | 0 dpf – 4 dpf | -0.126 | 0.115 | 127 | -1.103 | 0.5138 | -0.237 |
| sqrt(11-KT) | common nesting – white nesting | -0.749 | 0.732 | 125 | -1.023 | 0.9094 | -0.3051 |
| | common 0 dpf – white 0 dpf | -1.603 | 0.740 | 125 | -2.167 | 0.2607 | -0.6534 |
| | common 4 dpf – white 4 dpf | -5.036 | 0.749 | 125 | -6.726 | **<0.0001** | -2.0521 |
| | common nesting – common 0 dpf | 1.743 | 0.732 | 125 | 2.381 | 0.1708 | 0.7103 |
| | common nesting – common 4 dpf | 3.789 | 0.732 | 125 | 5.177 | **<0.0001** | 1.5440 |

| | | | | | | | |
|---|---|---|---|---|---|---|---|
| | common 0 dpf – common 4 dpf | 2.046 | 0.740 | 125 | 2.765 | 0.0700 | 0.8337 |
| | white nesting – white 0 dpf | 0.88 | 0.740 | 125 | 1.201 | 0.8358 | 0.3620 |
| | white nesting – white 4 dpf | -0.498 | 0.749 | 125 | -0.665 | 0.9854 | -0.2030 |
| | white 0 dpf – white 4 dpf | -1.387 | 0.749 | 125 | -1.852 | 0.4367 | -0.5650 |
| log(T) | common nesting – white nesting | -0.10345 | 0.178 | 125 | -0.580 | 0.9922 | -0.1729 |
| | common 0 dpf – white 0 dpf | -0.45584 | 0.180 | 125 | -2.528 | 0.1241 | -0.7621 |
| | common 4 dpf – white 4 dpf | -1.03637 | 0.183 | 125 | -5.678 | **<0.0001** | -1.7326 |
| | common nesting – common 0 dpf | 0.52757 | 0.178 | 125 | 2.957 | **0.0422** | 0.8820 |
| | common nesting – common 4 dpf | 0.93899 | 0.178 | 125 | 5.263 | **<0.0001** | 1.5698 |
| | common 0 dpf – common 4 dpf | 0.41142 | 0.180 | 125 | 2.281 | 0.2096 | 0.6878 |
| | white nesting – white 0 dpf | 0.17518 | 0.180 | 125 | 0.971 | 0.9262 | 0.2929 |
| | white nesting – white 4 dpf | 0.00607 | 0.183 | 125 | 0.033 | 1.0000 | 0.0102 |
| | white 0 dpf – white 4 dpf | -0.16911 | 0.183 | 125 | -0.927 | 0.9390 | -0.2827 |
| log(2-ME2) | common nesting – white nesting | -0.244 | 0.193 | 125 | -1.263 | 0.8046 | -0.377 |
| | common 0 dpf – white 0 dpf | -0.603 | 0.195 | 125 | -3.086 | **0.0294** | -0.931 |
| | common 4 dpf – white 4 dpf | -1.740 | 0.198 | 125 | -8.808 | **<0.0001** | -2.687 |
| | common nesting – common 0 dpf | 0.405 | 0.193 | 125 | 2.096 | 0.2963 | 0.625 |
| | common nesting – common 4 dpf | 1.333 | 0.193 | 125 | 6.900 | **<0.0001** | 2.058 |
| | common 0 dpf – common 4 dpf | 0.928 | 0.195 | 125 | 4.753 | **0.0001** | 1.433 |
| | white nesting – white 0 dpf | 0.046 | 0.195 | 125 | 0.235 | 0.9999 | 0.071 |
| | white nesting – white 4 dpf | -0.164 | 0.198 | 125 | -0.829 | 0.9617 | -0.253 |
| | white 0 dpf – white 4 dpf | -0.210 | 0.198 | 125 | -1.061 | 0.8956 | -0.324 |

**Supplementary Table 4.** Results of post-hoc tests and effect sizes corresponding to Table 2 (progestogens). Emmeans and effect sizes for pairwise differences in steroid levels are reported on the transformed (sqrt and log) scale. P-values are corrected for multiple testing using the tukey correction and effect size is Cohen's d. Significant p-values ($p<0.05$) are bolded.

| response variable | contrast | estimate | SE | df | t-ratio | p-value | effect size |
|---|---|---|---|---|---|---|---|
| sqrt(11β-OHP) | common nesting – white nesting | 0.6420 | 0.177 | 121 | 3.637 | **0.0053** | 1.085 |
| | common 0 dpf – white 0 dpf | 0.0716 | 0.185 | 121 | 0.386 | 0.9989 | 0.121 |
| | common 4 dpf – white 4 dpf | 0.6198 | 0.183 | 121 | 3.394 | **0.0117** | 1.047 |
| | common nesting – common 0 dpf | 1.0084 | 0.177 | 121 | 5.712 | **<0.0001** | 1.704 |
| | common nesting – common 4 dpf | 0.1559 | 0.179 | 121 | 0.872 | 0.9525 | 0.263 |
| | common 0 dpf – common 4 dpf | -0.8525 | 0.181 | 121 | -4.720 | **0.0001** | -1.440 |
| | white nesting – white 0 dpf | 0.4380 | 0.185 | 121 | 2.363 | 0.1779 | 0.740 |
| | white nesting – white 4 dpf | 0.1338 | 0.181 | 121 | 0.741 | 0.9764 | 0.226 |
| | white 0 dpf – white 4 dpf | -0.3042 | 0.188 | 121 | -1.622 | 0.5857 | -0.514 |
| log(pregnanolone) | common – white | 0.423 | 0.122 | 125 | 3.470 | **0.0007** | 0.609 |
| | nesting – 0 dpf | 0.807 | 0.149 | 125 | 5.409 | **<0.0001** | 1.161 |
| | nesting – 4 dpf | 0.134 | 0.148 | 125 | 0.906 | 0.6375 | 0.193 |
| | 0 dpf – 4 dpf | -0.672 | 0.151 | 125 | -4.456 | **0.0001** | -0.967 |

**Supplementary Table 5.** Results of post-hoc tests and effect sizes corresponding to Table 3 (corticosteroids). Emmeans and effect sizes for pairwise differences in steroid levels are reported on the transformed (log) scale. P-values are corrected for multiple testing using the tukey correction and effect size is Cohen's d. Significant p-values ($p<0.05$) are bolded.

| response variable | contrast | estimate | SE | df | t-ratio | p-value | effect size |
|---|---|---|---|---|---|---|---|
| log(cortisol) | nesting – 0 dpf | -0.692 | 0.251 | 127 | -2.752 | **0.0185** | -0.584 |
| | nesting – 4 dpf | 0.483 | 0.253 | 127 | 1.909 | 0.1404 | 0.407 |
| | 0 dpf – 4 dpf | 1.174 | 0.254 | 127 | 4.620 | **<0.0001** | 0.991 |
| log(cortisone) | nesting – 0 dpf | -0.378 | 0.203 | 127 | -1.865 | 0.1532 | -0.395 |
| | nesting – 4 dpf | 0.260 | 0.204 | 127 | 1.272 | 0.4135 | 0.272 |
| | 0 dpf – 4 dpf | 0.637 | 0.205 | 127 | 3.110 | **0.0065** | 0.667 |
| log(aldosterone) | nesting – 0 dpf | -0.320 | 0.159 | 127 | -2.017 | 0.1122 | -0.428 |
| | nesting – 4 dpf | 0.151 | 0.160 | 127 | 0.946 | 0.6124 | 0.202 |
| | 0 dpf – 4 dpf | 0.471 | 0.161 | 127 | 2.935 | **0.0110** | 0.630 |

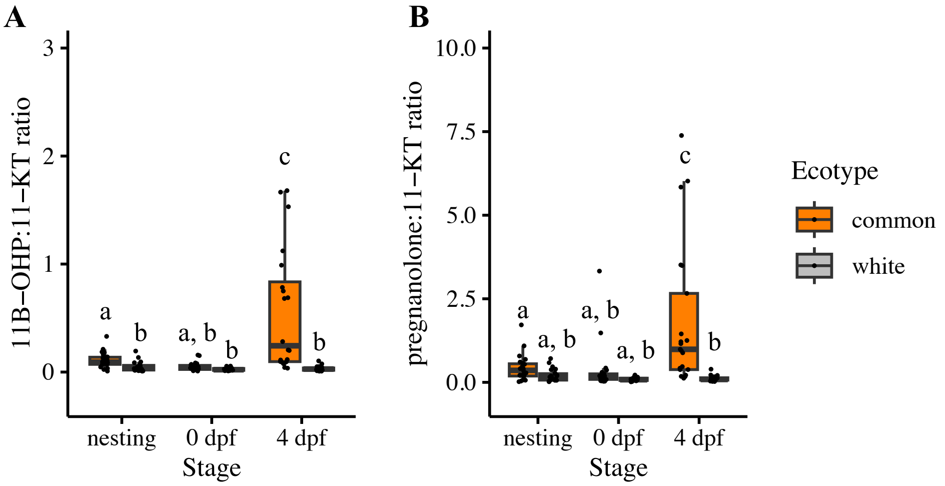


**Supplementary Figure 1.** *The ratio of progestogen metabolites to 11-KT, the major androgen in teleosts, was low across all groups except parenting commons (4 dpf). (A)* 11β-OHP to 11-KT ratio (11β-OHP levels divided by 11-KT levels for each male). *(B)* Pregnanolone to 11-KT ratio (11β-OHP levels divided by 11-KT levels for each male). Boxplots show the median (thick black bar), the first and third quartiles (bottom and top edges of box), and 1.5 times the interquartile range (whiskers). Commons are shown in orange, and whites are shown in grey. Each point represents an individual male. Different letters denote significant differences ($p<0.05$) between groups in estimated marginal means (EMM) post-hoc tests after Tukey correction. Sample sizes for 11β-OHP are as follows: n=23 nesting commons, n=22 nesting whites, n=22 0 dpf commons, n=19 0 dpf whites, n=21 4 dpf commons, and n=21 4 dpf whites. Samples sizes were the same for pregnanolone except for 0 dpf whites (n=20) and 4 dpf commons (n=22). One common (4 dpf) and one white (0 dpf) had unusually high ratios due to very low values for 11-KT; these points have been omitted to make between-group differences easier to visualize.

**Supplementary Table 6**. Results from linear mixed models testing for effects of ecotype and reproductive stage on the ratio of progestogens to 11-KT. Hormone levels were log transformed before analysis to better fit the assumptions of normality, and models included a random effect of year to account for the fact that data were collected over two years. Significant p-values ($p<0.05$) are bolded. Degrees of freedom (numerator, denominator) were estimated using the Satterthwaite method.

| **Steroid** | ***F*** | **df** | **P** |
|---|---|---|---|
| **11β-OH-Progesterone (11β-OHP): 11-KT** | | | |
| *ecotype* | 43.910 | 1, 122 | **<0.001** |
| *stage* | 9.167 | 2, 122 | **<0.001** |
| *ecotype*stage* | 14.235 | 2, 122 | **<0.001** |
| **Pregnanolone: 11-KT** | | | |
| *ecotype* | 34.803 | 1, 123.00 | **<0.001** |
| *stage* | 6.422 | 2, 123.01 | **0.002** |
| *ecotype*stage* | 10.274 | 2, 123.01 | **<0.001** |

**Supplementary Table 7.** Results of post-hoc tests and effect sizes corresponding to Supplementary Table 6 (progestogen:androgen ratios). Emmeans and effect sizes for pairwise differences in steroid levels are reported on the transformed (sqrt and log) scale. P-values are corrected for multiple testing using the tukey correction and effect size is Cohen's d. Significant p-values (p<0.05) are bolded.

| response variable | contrast | estimate | SE | df | t-ratio | p-value | effect size |
|---|---|---|---|---|---|---|---|
| log(11β-OHP:11-KT) | common nesting – white nesting | 0.9156 | 0.316 | 121 | 2.898 | **0.0497** | 0.8646 |
| | common 0 dpf – white 0 dpf | 0.2013 | 0.332 | 121 | 0.607 | 0.9904 | 0.1900 |
| | common 4 dpf – white 4 dpf | 2.6112 | 0.327 | 121 | 7.990 | **<0.0001** | 2.4658 |
| | common nesting – common 0 dpf | 0.8056 | 0.316 | 121 | 2.550 | 0.1181 | 0.7608 |
| | common nesting – common 4 dpf | -1.3940 | 0.320 | 121 | -4.354 | **0.0004** | -1.3164 |
| | common 0 dpf – common 4 dpf | -2.1996 | 0.323 | 121 | -6.805 | **<0.0001** | -2.0771 |
| | white nesting – white 0 dpf | 0.0912 | 0.332 | 121 | 0.275 | 0.9998 | 0.0862 |
| | white nesting – white 4 dpf | 0.3016 | 0.323 | 121 | 0.933 | 0.9372 | 0.2848 |
| | white 0 dpf – white 4 dpf | 0.2104 | 0.336 | 121 | 0.627 | 0.9888 | 0.1987 |
| log(pregnanolone:11-KT) | common nesting – white nesting | 0.668 | 0.351 | 123 | 1.907 | 0.4030 | 0.569 |
| | common 0 dpf – white 0 dpf | 0.445 | 0.363 | 123 | 1.225 | 0.8236 | 0.379 |
| | common 4 dpf – white 4 dpf | 2.539 | 0.359 | 123 | 7.079 | **<0.0001** | 2.160 |
| | common nesting – common 0 dpf | 0.467 | 0.351 | 123 | 1.332 | 0.7664 | 0.397 |
| | common nesting – common 4 dpf | -1.486 | 0.351 | 123 | -4.240 | **0.0006** | -1.265 |
| | common 0 dpf – common 4 dpf | -1.953 | 0.354 | 123 | -5.512 | **<0.0001** | -1.662 |
| | white nesting – white 0 dpf | 0.244 | 0.363 | 123 | 0.671 | 0.9848 | 0.207 |
| | white nesting – white 4 dpf | 0.384 | 0.359 | 123 | 1.071 | 0.8921 | 0.327 |
| | white 0 dpf – white 4 dpf | 0.140 | 0.368 | 123 | 0.381 | 0.9989 | 0.119 |

**Supplementary Table 8.** Results of post-hoc tests and effect sizes for gluing behavior. Emmeans and effect sizes for pairwise differences are reported on the log scale because gluing data were analyzed with a negative binomial generalized linear mixed model. P-values are corrected for multiple testing using the tukey correction and effect size is Cohen's d. Significant p-values ($p<0.05$) are bolded. We focus on comparisons at 0 and 4 dpf.

| response variable | contrast | estimate | SE | df | z-ratio | p-value | effect size |
|---|---|---|---|---|---|---|---|
| number of glues | common 0 dpf – common 4 dpf | 2.23072 | 0.396 | Inf | 5.630 | **<0.0001** | 2.23072 |
| | white 0 dpf – white 4 dpf | 0.37596 | 0.326 | Inf | 1.154 | 0.9789 | 0.37596 |
| | common 0 dpf – white 0 dpf | 0.53712 | 0.343 | Inf | 1.564 | 0.8656 | 0.53712 |
| | common 4 dpf – white 4 dpf | -1.31764 | 0.442 | Inf | -2.980 | 0.0850 | -1.31764 |
